# LACK OF OXYGEN AND/OR GLUCOSE DIFFERENTIALLY POTENTIATES Aβ40_E22Q_- AND Aβ42-INDUCED CEREBRAL ENDOTHELIAL CELL DEATH, BARRIER DYSFUNCTION AND ANGIOGENESIS IMPAIRMENT

**DOI:** 10.1101/2025.10.05.680546

**Authors:** Ashley Carey, Tetyana Buzhdygan, Silvia Fossati

## Abstract

**Background:** Disrupted brain hemodynamics and cerebrovascular damage resulting in cerebral hypoperfusion occur early within Alzheimer’s Disease (AD) pathogenesis. Cerebral hypoperfusion is also an extremely common consequence of cardiovascular risk factors and diseases (CVRFs/CVDs), which usually manifest in midlife, when AD pathology initiates, and actively contribute to AD onset and progression. Previously our lab has demonstrated that the vasculotropic Dutch mutant, AβQ22, and Aβ42 promote endothelial cells (ECs) apoptosis, barrier permeability, and angiogenic impairments. Prior research has indicated that hypoperfusion promotes analogous EC dysfunction. Aβ deposition occurs within a hypoperfused environment in AD, but whether exposure of cerebral ECs to Aβ under hypoperfusion results in potentiated cerebral EC dysfunction through activation of common molecular mechanisms remained unknown.

**Methods:** Human cerebral ECs were treated with Aβ40-Q22 or Aβ42, glucose deprivation (GD), or a combination of both, under normoxia or hypoxia conditions. Cell death mechanisms (apoptosis/necrosis), endothelial barrier dysfunction/permeability (TEER/barrier-regulating proteins/proinflammatory activation), and angiogenesis impairment (vessel branching/VEGF signaling) were evaluated.

**Results:** Reduction of glucose and/or oxygen potentiates Aβ-induced cerebral EC death, barrier instability, junction protein dysregulation, inflammatory activation, and angiogenesis/wound healing failure. In particular, hypoperfusion exacerbates AβQ22-mediated cerebral EC apoptosis, TEER/ZO1 decreases, ICAM1, IL6, and IL8 upregulation, monocyte migration, and wound healing impairments. Differentially, when in combination with Aβ42, hypoperfusion more strongly potentiates cerebral EC necrosis as well as increases in MMP2, phosphorylated claudin-5, IFNγ, and IL12p70 expression. Additionally, this study identified that GD exerts stronger effects on promoting increases in cerebral EC caspase-3 activation, apoptosis, and MMP2/ICAM1 expression, while hypoxia particularly increases necrosis, ZO1 expression, and pro-angiogenic protein expression.

**Conclusions:** This study reveals specific and selective mechanisms through which hypoxia, low glucose and amyloidosis mutually operate to produce brain EC dysfunction and death, highlighting new potential molecular targets against vascular pathology in AD/CAA comorbid with hypoperfusion conditions.

**Highlights:**

- Depriving cerebral endothelial cells of glucose and/or oxygen potentiates Aβ-induced endothelial dysfunction, differentially promoting increased cell death, barrier instability and dysregulation of blood brain barrier proteins, inflammatory activation, and angiogenesis and wound healing failure, in relation to the specific peptide and low glucose or oxygen conditions.
- Under hypoperfusion conditions, AβQ22 more strongly exacerbates increases in apoptosis, ICAM1, IL6, and IL8 expression, and monocyte migration and decreases in TEER, ZO1 expression, and wound healing, revealing that the vasculotropic AβQ22 produces even stronger vascular effects when in combination with hypoperfusion.
- Under hypoperfusion conditions, Aβ42 more strongly potentiates increases in necrosis and MMP2, phosphorylated claudin-5, IFNγ, and IL12p70 expression.
- Glucose deprivation exerts stronger effects on increasing caspase-3 activation, apoptosis, and MMP2 and ICAM1 expression, while hypoxia displays stronger effects on increasing necrosis and ZO1 and pro-angiogenic protein expression.
- We demonstrated that AβQ22 more intensely promotes vascular dysfunction when in combination with hypoperfusion conditions versus Aβ42.
- verall, results from this study point to the importance of monitoring and preventing cerebral hypoperfusion particularly during midlife, when AD pathology begins to develop, to prevent this early pathology from working with Aβ to create a more detrimental dementia trajectory, and highlights new targets for possible therapeutic or preventive strategies.

## Introduction

Alzheimer’s Disease (AD) is the most prevalent age-associated dementia and is currently affecting more than 6.9 million Americans, with case numbers rising exponentially and the annual death rate climbing [1]. The deposition of amyloid beta (Aβ) plaques, accumulation of hyperphosphorylated tau neurofibrillary tangles, chronic neuroinflammation and cerebrovascular dysfunction pathologically define AD, with these pathologies promoting neuronal death and ultimately cognitive decline [2]. A common phenomenon within the general aging population and particularly in 85-95% of AD patients is Cerebral Amyloid Angiopathy (CAA), characterized by Aβ deposition within and around the cerebral vasculature [2–4]. CAA’s consequences include focal ischemia and cerebral hypoperfusion (CH), cerebral blood flow (CBF) impairments, blood brain barrier (BBB) dysfunction, as well as cerebral hemorrhages and microinfarcts [2, 5]. CAA deposits are mainly composed of the Aβ40 species, which tend to accumulate within the cerebral vessel walls, while plaques mostly containing Aβ42 are located within the cerebral parenchyma [5, 6]. The vasculotropic familial Dutch mutant, Aβ40_E22Q_ (AβQ22), contains an amino acid 22 glutamic acid to glutamine substitution that fast-tracks the peptides’ oligomerization. Specifically, this Aβ mutant is closely associated with early onset CAA, with disease progression beginning as early as 30-40 years of life and pathologies encompassing hemorrhages, strokes, and dementia [4, 7]. AβQ22 is more aggregation prone and exerts more aggressive effects on endothelial cells (ECs) compared to wild-type (WT) Aβ40 [4], thus utilizing AβQ22 within our *in vitro* experiments serves as a vital tool to investigate the specific effects of vascular Aβ.

It is now widely recognized that more than half of dementia cases are mixed pathology dementias, with AD and cerebrovascular pathology being the most frequent combination [8]. Additionally, mixed vascular and Alzheimer’s dementia is common within the general aging population, with a prevalence of 22% [9]. Mounting evidence over the past decades reveals that cerebrovascular dysfunction and CBF impairments, resulting in CH, are likely strong influencers of early AD pathogenesis, occurring prior to significant amyloid accumulation [10–13]. Recent studies have proposed that chronic CH is a driver of neurodegeneration by promoting cerebral oxygen and nutrient deficits as well as increased oxidative stress and neuroinflammation, resulting in neuronal death and accelerated cognitive decline [14, 15]. Studies have revealed not only that AD patients demonstrate CH in distinct cortical areas [16–18], but within the general population, increased CH has also been predictive of exacerbated dementia risk and cognitive decline [19–21]. Taken together, it is clear there is a vital necessity to better understand the contributions of early cerebrovascular changes such as CH to the mechanisms responsible for the vascular contributions to cognitive impairment and dementia (VCID), as these mechanisms could be targeted for prevention and treatment options.

A majority of cardiovascular diseases and risk factors have been found to promote CH [22] and to increase dementia and AD risk [15]. The natural aging process itself is also known to promote CH, with studies demonstrating a 20% decrease in CBF by age 60 [22], resulting in decreases in oxygen and nutrient availability, and potentiated neuronal death [22]. Notably, hypertension has been found to promote CH, with the proposed pathological mechanisms involving cerebral ischemia and hemorrhages, capillary rarefaction, endothelial dysfunction and death, BBB permeability and neuroinflammation [23]. Similarly, ischemic strokes are known to promote CH, as they specifically result from vessel occlusions caused by vascular complications (blood clots, stenosis, intracranial atherosclerotic plaques) [24]. Additionally, CH and Aβ have been shown to produce similar cerebrovascular pathologies. Despite this, whether CH and amyloidosis operate through common molecular mechanisms to promote cerebral endothelial dysfunction and whether they do so in an additive or synergistic manner remains unknown. Dissecting the molecular pathways activated by both Aβ and CH that result in endothelial pathology will be imperative for pinpointing novel molecular targets that could be utilized to treat the comorbid vascular effects of Aβ and CH within AD and dementia patients and the aging population.

Our lab was pioneer in outlining the mechanisms by which Aβ, particularly AβQ22 and Aβ42, affect cerebral EC function. We have demonstrated that treatment of human cerebral microvascular ECs (HCMECs) with AβQ22 and Aβ42 results in increased mitochondria-mediated apoptosis [4, 25, 26]. Additionally, our studies have revealed that AβQ22 and Aβ42 treatments cause HCMEC barrier dysfunction, promoting trans-endothelial electrical resistance (TEER) loss and dysregulation of BBB-regulating proteins [26–28], and also that these Aβ species decrease HCMEC angiogenic and wound healing capabilities [26, 28]. Likewise, previous literature reveals that oxygen glucose deprivation (OGD), an *in vitro* model of hypoperfusion, promotes similar cerebral EC dysfunction, specifically promoting apoptosis and necrosis [29], barrier permeability [30, 31], and angiogenesis/wound healing impairments [32–35]. This study aims to determine whether partial OGD potentiates specific mechanisms of Aβ-induced HCMEC death, barrier dysfunction and angiogenic impairments. An additional goal of this study is to understand whether the EC dysfunction resulting from the combination of OGD and Aβ is Aβ-species specific. We hypothesize that OGD will exacerbate Aβ-induced HCMEC apoptosis, barrier impairments, and angiogenic deficits in an additive manner and through activation of common molecular pathways, thus fast-tracking cerebrovascular pathological progression and increasing dementia risk.

## Material and Methods

### Cell Culture

HCMECs/D3, immortalized human cerebral microvascular ECs, were acquired from Babette Weksler (Cornell University) [4]. EBM-2 (Lonza) supplemented with growth factors (Hydrocortisone, hFGF-B, VEGF, R3-IGF-1, ascorbic acid, hEGF, GA-1000) and 5% fetal bovine serum (FBS) was used to grow the cells. Cells were maintained in a humidified cell culture incubator (37°C) under a 5% CO_2_ atmosphere. The EVOS M5000 Imaging System (ThermoFisher) was utilized for cell visualization and imaging.

### AβQ22 and Aβ42 Peptide

The Aβ42 peptide, the most aggregation prone brain amyloid, and the genetic variant of the WT Aβ40 peptide containing the E22Q vasculotropic substitution (AβQ22), were utilized for cell treatments. Specifically, AβQ22 is the synthetic homolog of the amyloid subunit present in the vascular deposits in sporadic and familial Dutch-AD cases. Peptide synthesis was performed by Peptide 2.0. Aβ42 and AβQ22 were dissolved to 1mM in 1,1,1,3,3,3-hexafluoro-2-propanol (HFIP; Sigma) and incubated for 24h to breakdown pre-existing β-sheet structures. After incubation, the peptide was lyophilized and then dissolved to a 10mM concentration in DMSO, followed by the addition of dH_2_O to a 1mM concentration. Aβ peptides were further diluted in culture media (DMEM (Gibco), 1% FBS and no growth factors) to the required experimental concentrations.

### Oxygen and Glucose Deprivation Treatments

Glucose-free DMEM (Gibco) was supplemented with 1M glucose (Agilent) to achieve final concentrations of 0.1mg/ml, defined as glucose deprivation (GD), and 1mg/ml, defined as normal glucose. Partial oxygen deprivation (hypoxia) was achieved by placing cells in a hypoxic chamber (Coy Laboratories), maintained at 1% oxygen and 5% CO_2_. OGD was achieved by treating cells with GD media and hypoxia (as above).

### Cell Death ELISA

Apoptosis was evaluated by measuring fragmented nucleosome levels using the Cell Death Detection ELISA^Plus^ (Roche) according to the manufacturer’s instructions. HCMECs were seeded and, following 24h, treated with 25µM AβQ22 or 5µM Aβ42 (concentrations that we know from our previous studies cause EC apoptosis) [25, 28], GD, or a combination of Aβ and GD under conditions of normoxia, hypoxia, or hypoxia/reoxygenation (HR = 24h hypoxia, followed by 24h normoxia) for 48h. Extranuclear DNA-histone fragmented complexes were measured as absorbance using the SpectraMax i3x Multi-Mode Microplate Reader (Molecular Devices). Results are expressed as percent change versus untreated controls.

### Lactate Dehydrogenase Assay

Necrosis was assessed by measuring lactate dehydrogenase (LDH) levels released within cell media utilizing a LDH assay kit (Roche) according to the manufacturer’s instructions. HCMECs were seeded for 24h and subsequently treated with 25µM AβQ22 or 5µM Aβ42, GD, or a combination of both under conditions of normoxia, hypoxia, or HR for 48h. Cell media was collected and LDH levels were measured as absorbance (492 nm) using the SpectraMax i3x Microplate Reader. Results are expressed as percent change versus untreated controls.

### Cell Event Fluorescent Caspase 3/7 Assay

Cleaved/active caspase-3/7 expression was evaluated with the CellEvent^TM^ Caspase-3/7 Green Detection Reagent (ThermoFischer). Cells were seeded in a 96-well plate (10,000 cells/well) and treated for 24h with 25µM AβQ22 or 5µM Aβ42, GD, or a combination of both under normoxia or hypoxia conditions. Post-treatment, cell media was removed, the fluorescent Caspase-3/7 Detection Reagent was added to each well (5µM), and cells were incubated for 1h at 37°C. Cells were imaged with bright-field and GFP filters via the EVOS M5000 imaging system (4 randomized images/well). Percent of cleaved caspase-3/7 positive cells was determined by counting the total cell number in each picture, followed by the creation of a mask for the fluorescent signal to count the amount of cells demonstrating active caspase-3/7 fluorescence [% Caspase-3/7+ Cells = (# of cells with cleaved caspase-3/7**/**total cell #)*100].

### ECIS Trans-Endothelial Electrical Resistance

The ECIS Zθ system (Applied Biophysics) was utilized to assess HCMEC barrier integrity. Experimental procedures were performed in 8-well ECIS (8WE10+) 40 electrode gold-plated arrays, which were pre-treated according to the manufacturer’s recommendations. HCMECs were seeded and monitored for 48h until TEER reached a plateau at a frequency of 4000 Hz, which is indicative of complete barrier formation. The cell monolayers were then treated with 25µM AβQ22 or 5µM Aβ42, GD, or a combination of both under normoxia or hypoxia and followed for 24h post-treatment. Barrier permeability was defined as a decrease in TEER at 4000 Hz versus untreated controls.

### ECIS Wound Healing Assay

HCMEC wound healing capability was evaluated utilizing the corresponding ECIS Zθ assay. Experimental procedures were performed in 8-well ECIS (8W1E) gold-plated arrays containing one central electrode, and the arrays were pre-treated according to the manufacturer’s recommendations. HCMECs were seeded and monitored for 48h until TEER reached a plateau at a 4000 Hz frequency (barrier formation). The cell monolayers were then treated with 25µM AβQ22 or 5µM Aβ42, GD, or a combination of both under normoxia or hypoxia conditions. One hour post-treatment, a wound was inflicted to the EC monolayer lasting 20 seconds (60,000 Hertz; amplitude=5V; wound current=1400µAmps) and wound healing was assessed for 48h post-wound. Impairments in wound healing were quantified as decreased post-wound recovery TEER versus untreated controls.

### Western Blot

Evaluation of ZO1 (Invitrogen; 61-7300), ICAM (Invitrogen; MA5407), phosphorylated claudin-5 (abcam; ab172968), MMP2 (abcam; ab86607), phosphorylated VEGFR2 Y1175 (abcam; ab194806), and VEGF-A (Proteintech; 19003-1-AP) expression was performed using western blot (WB) analysis after electrophoretic separation on 4-12% Bolt Bis-Tris SDS polyacrylamide gels. Anti-β actin (Millipore; MAB1501) was utilized for normalization. Proteins were electrotransferred to nitrocellulose membranes (0.45μm pore; Cytiva LifeSciences) at 110 V for 70 min, using towbin buffer, containing 20% (v/v) methanol. Intercept Blocking Buffer (Licor) was utilized to block membranes and membranes were subsequently immunoreacted with the corresponding primary antibodies for each experiment, followed by incubation with the appropriate anti-rabbit or anti-mouse secondary antibodies (1/20,000; Licor). Membranes were developed utilizing the LICOR Odyssey CLx Immunoblot Imager and blots were analyzed with the LICOR Image Studio software.

### Proinflammatory Panel MSD

Proinflammatory cytokine expression was measured via the V-PLEX proinflammatory panel 1 kit from MesoScale Discovery (MSD), which measures expression levels of 10 cytokines/sample (IL1β, IL2, IL4, IL6, IL8, IL10, IL12p70, IL13, IFNγ, TNFα). HCMECs were seeded for 24h and then treated with 25µM AβQ22 or 5µM Aβ42, GD, or a combination of both under normoxia or hypoxia conditions for 6h. Media was collected post-treatment and stored at -80°C until the MSD was conducted. The assay was performed in accordance with the manufacturer’s recommendations and sample protein concentration was utilized for normalization.

### U937 Human Monocytes

The human monocyte cell line U937 was obtained from ATCC and maintained in RPMI-1640 media supplemented with 10% heat-inactivated pooled FBS. Cells were serum-starved for 24h prior to adhesion and extravasation assays.

### Monocyte Trans-endothelial Extravasation Assay

A monocyte extravasation assay was performed as previously described [36]. Briefly, HCMECs were plated on rat tail collagen type I-coated FluoroBlok^™^ HTS 96-well inserts with 8µM pores (Corning) (2.5×10^4^ cells/insert). After confluent monolayer formation, HCMECs were treated with 25µM AβQ22 or 5µM Aβ42, OGD, or a combination of the two for 24h. All conditions were also supplemented with 1× GlutaMAX. On extravasation challenge day, U937 monocytes were loaded with Calcein-AM at 5µM/1×10^6^ cells for 45 min. Permeable supports with HCMEC monolayers were rinsed post-treatment and transferred to wells filled with phenol-free DMEM supplemented with 1g/L glucose, 1x GlutaMAX, and 10% FBS. Calcein-AM labeled U937 (40,000 cells/support) were added to the luminal side of the HCMECs and allowed to migrate and adhere to the abluminal side of the supports for 6h at 37°C. For fluorescence measurement, supports were transferred to clean 24-well plates filled with HBSS and fluorescence was acquired with a Spectramax M5 fluorescence plate reader. The data was calculated based on the standard curve derived from fluorescent intensity of known numbers of labeled monocytes.

### Angiogenesis Inhibition Assay

Angiogenic potential was evaluated with the Millicell μ-Angiogenesis Inhibition Assay (Millipore) according to the manufacturer’s instructions. HCMECs were seeded in μ-angiogenesis slides containing ECMatrix gel solution and simultaneously treated with a sublethal concentration (1µM) of AβQ22 or Aβ42, GD, or a combination of both under normoxia or hypoxia conditions [28]. Tube formation was assessed after 6h by acquiring pictures with the EVOS M5000 imaging system (4 randomized images/well). Capillary branches meeting length criteria were counted within the images.

### HIF1α ELISA

HIF1α expression was measured with a Total HIF1α Duoset ELISA (Bio-techne R&D Systems), following the manufacturer’s recommendations. HCMECs were seeded for 24h and then treated with 25µM AβQ22 or 5µM Aβ42, GD, or a combination of both under normoxia or hypoxia conditions for 6h. Post-treatment, protein lysate was obtained and stored at -80°C until the ELISA was conducted. Sample protein concentration was utilized for normalization.

### VEGF-A ELISA

Soluble VEGF-A release was measured via the VEGF-A Quantikine ELISA (Bio-techne R&D Systems), following the manufacturer’s recommendations. HCMECs were seeded for 24h and then treated with 25µM AβQ22 or 5µM Aβ42, GD, or a combination of both under normoxia or hypoxia conditions for 6h and 24h. Post-treatment, media was collected and stored at -80°C until the ELISA was conducted. Sample protein concentration was utilized for normalization.

### Statistical Analysis

All experimental graphs are representative of at least 3 independent experiments with 2 or more technical replicates. All data is represented as means±SEM. Statistical significance for ECIS TEER and wound healing was evaluated by one-way ANOVA followed by Tukey’s post-hoc test and statistical significance for all other experiments was assessed by two-way ANOVA followed by Tukey’s post-hoc test using GraphPad Prism 9. Statistically significant differences required a p-value≤0.05.

## Results

### Exposure of HCMECs to AβQ22 or Aβ42 and GD, under varying oxygen conditions, differentially affects apoptosis and necrosis

It has been shown that the vasculotropic Aβ40 variant, AβQ22, and the aggregation-prone Aβ42 peptide promote increased fragmented nucleosomes formation, indicative of elevated apoptosis, as well as increased LDH release, indicative of increased necrosis levels [4, 26, 37]. OGD, a widely used *in vitro* model of CH, has also been found to promote increased brain microvascular EC death, specifically through the apoptotic and necrotic pathways [29]. Aβ accumulation and hypoperfusion are simultaneously occurring within AD and CAA brains [38, 39] and both pathological factors can promote cerebrovascular cell death. However, it is currently unknown whether the combined challenge of specific Aβ species and OGD will exacerbate cerebral EC death through common cell death mechanisms, specifically apoptosis and necrosis. To investigate this, we measured cleaved caspase-3, apoptosis, and necrosis levels in HCMECs treated with 25µM AβQ22 or 5µM Aβ42 (concentrations that we know start activating apoptotic pathways [25, 26, 28]), GD, or a combination of both under varying oxygenation conditions. The rationale for treating HCMECs with lower concentrations of Aβ42 versus AβQ22 was based on our lab’s prior findings demonstrating that Aβ42’s toxic effects are more rapid and drastic versus Aβ40 and its variants, due to Aβ42 being extremely aggregation prone and showing faster oligomerization kinetics [4, 26]. Due to this, HCMECs were treated with a higher concentration of AβQ22 (25µM) versus Aβ42 (5µM) to take into consideration these aggregation and toxicity differences. CH conditions within this study were modeled by exposing HCMECs to OGD conditions, defined here as partial depletion of oxygen (hypoxia: 1% O_2_) and glucose (controls: 1mg/ml vs. GD: 0.1mg/ml).

Cleaved (activated) caspase-3/7, the executioner caspases within the apoptotic pathway, were measured using a fluorescent active caspase-3/7 assay following 24h treatment. HCMECs exposed to AβQ22 or Aβ42 and GD both demonstrated a potentiated increase in cleaved caspase-3/7 levels compared to control cells and the challenges alone, with Aβ42+GD more significantly increasing cleaved caspase-3/7 levels versus AβQ22+GD exposed cells (**Fig. 1A)**. HCMECs treated with Aβ and/or GD under hypoxic conditions demonstrated a similar trend of potentiated cleaved caspase-3 levels, although hypoxia did not further exacerbate cleaved caspase-3 increases when added to GD (OGD) (**Fig. 1A**). To confirm that caspase-3 activation resulted in the execution of the apoptotic pathway, we measured DNA fragmentation, indicative of the terminal stage of apoptosis, after 48h challenge. Treatment of HCMECs with GD significantly potentiated AβQ22-mediated increases in apoptosis versus controls and each challenge alone (**Fig. 1B**). However, we did not observe this increase with Aβ42+GD (**Fig. 1C**), as Aβ42 alone already resulted in much higher apoptosis levels versus AβQ22. Instead, Aβ42+GD challenge resulted in a significant increase in necrosis (**Fig. 1C**). Interestingly, hypoxia and GD combined challenge (OGD), resulted in a significant increase in Aβ42-induced necrosis, while reducing the levels of apoptosis caused by this peptide. Since a common phenomenon in ischemic or hypoxic tissue environments is hypoxia-reperfusion injury, where blood flow restoration may cause additional damage, resulting in exacerbated EC death [40], a HR condition (24h hypoxia, then 24h normoxia) was also considered when measuring apoptosis and necrosis. HR itself significantly increased HCMEC apoptosis levels (**Fig. 1B/C**), while hypoxia and HR did not further potentiate HCMEC apoptosis levels when in combination with Aβ or GD. Aβ42 and OGD exposure resulted in an additive increase in HCMEC necrosis levels (**Fig. 1C**). HR alone also significantly increased HCMEC necrosis (**Fig. 1B/C**), with ECs exposed to HR+AβQ22 or HR+AβQ22+GD demonstrating the most significant increase in necrosis versus controls (**Fig. 1B**).

**Figure 1.**
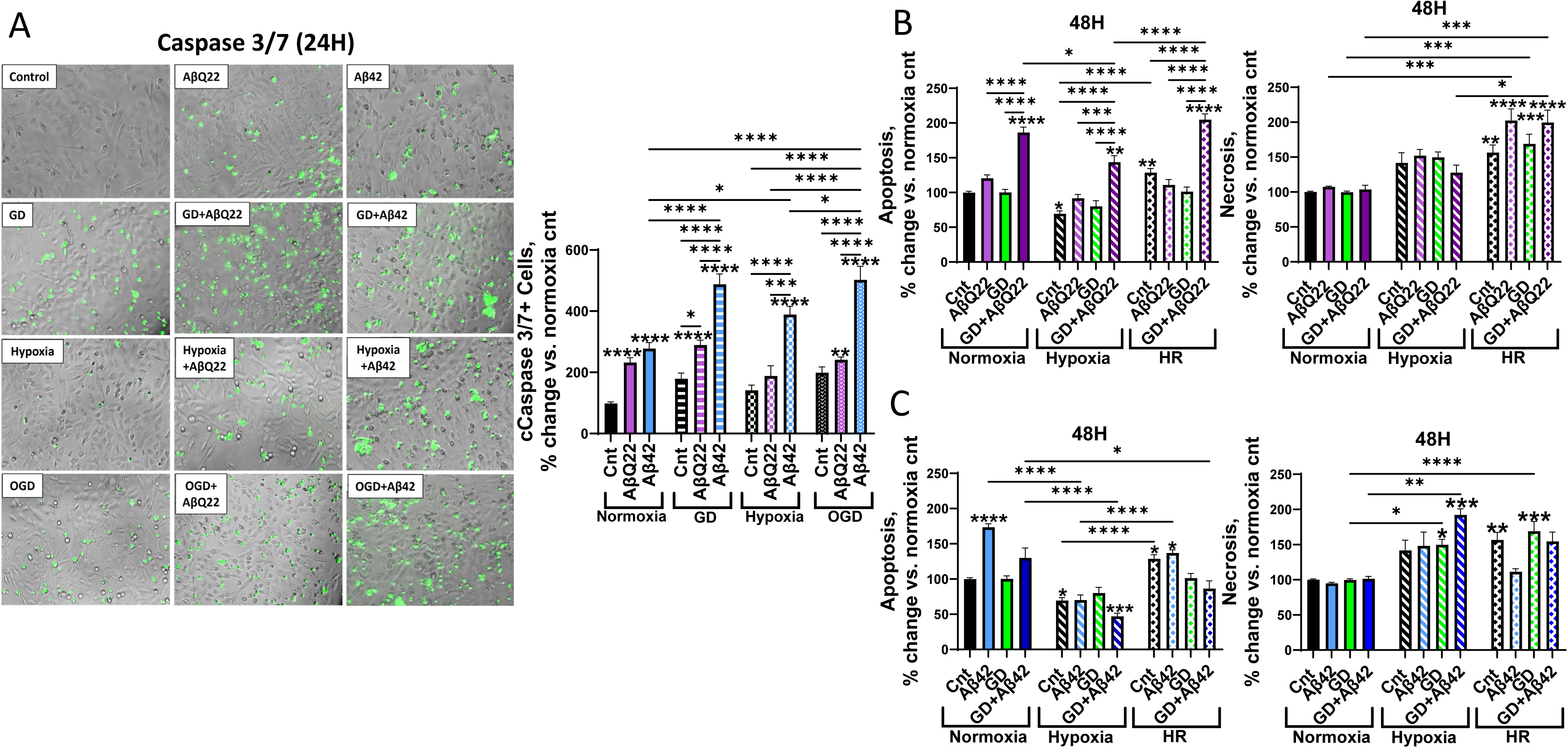
AβQ22 and Aβ42 in the presence of GD or OGD differentially affect apoptosis and necrosis in HCMECs. **(A)** HCMECs were treated with 25µM AβQ22 or 5µM Aβ42, GD, or a combination of the two for 24h under conditions of normoxia or hypoxia. The amount of cleaved caspase 3/7 present in the cells was measured with ThermoFisher’s Cell Event Caspase 3/7 fluorescence assay (Invitrogen’s EVOS M5000 microscope for imaging). Percent of cleaved caspase 3/7+ cells was calculated as (caspase 3/7+ cells/total cell number)*100. Data is represented as % change vs. normoxia control. N=3 experiments with 2 or more technical replicates; two-way ANOVA, Tukey’s post-test. **(B/C)** HCMECs were treated with 25µM AβQ22 **(B)** or 5µM Aβ42 **(C)**, GD, or a combination of the two for 48h under conditions of normoxia, hypoxia, or HR. Apoptosis levels were assessed by measuring DNA fragmentation utilizing the Cell Death ELISA^PLUS^ assay (Roche). Necrosis levels were assessed by measuring LDH in cell media with a LDH assay. Data is represented as % change vs. normoxia control. N≥3 experiments with 2 or more technical replicates; two-way ANOVA, Tukey’s post-test. (**** p<0.0001, ***p<0.001, **p<0.01, *p<0.05).

### Treatment of HCMECs with AβQ22 and OGD additively decreases TEER and wound healing capabilities

Our lab’s prior findings have demonstrated that AβQ22 and Aβ42 promote HCMEC barrier permeability, reducing the electrical resistance between ECs [26, 28]. Additionally, our recent study has shown that AβQ22 impairs HCMEC wound healing capabilities [28]. OGD has also been shown to promote barrier dysfunction and permeability [30, 31, 41] as well as wound healing impairments [32, 33] in cerebral ECs. To determine whether HCMECs exposed to either Aβ species and/or OGD demonstrate additive increases in endothelial barrier permeability and wound healing impairments, ECIS technology was utilized. Once a stable EC barrier was established, HCMECs were treated with AβQ22 (25µM) or Aβ42 (5µM), GD, or a combination of both under normoxia or hypoxia and TEER was measured over time for 24h. As expected, AβQ22 and Aβ42 significantly decreased HCMEC TEER versus controls (**Fig. 2A/B**). HCMECs exposed to AβQ22+OGD demonstrated an additive decrease in TEER compared to the treatments alone (**Fig. 2A**). Similarly, when Aβ42 was given in combination with either GD, hypoxia, or OGD, HCMEC TEER was more significantly reduced compared to cells treated with Aβ42 alone (**Fig. 2B**), but Aβ42 addition, at this concentration, did not have a significant additive effect on OGD-induced TEER decreases at later timepoints, possibly due to a floor effect where maximal TEER decreases were achieved, suggesting that the barrier impairment in Aβ42+OGD conditions is primarily driven by OGD, rather than Aβ42 (**Fig. 2B**).

**Figure 2.**
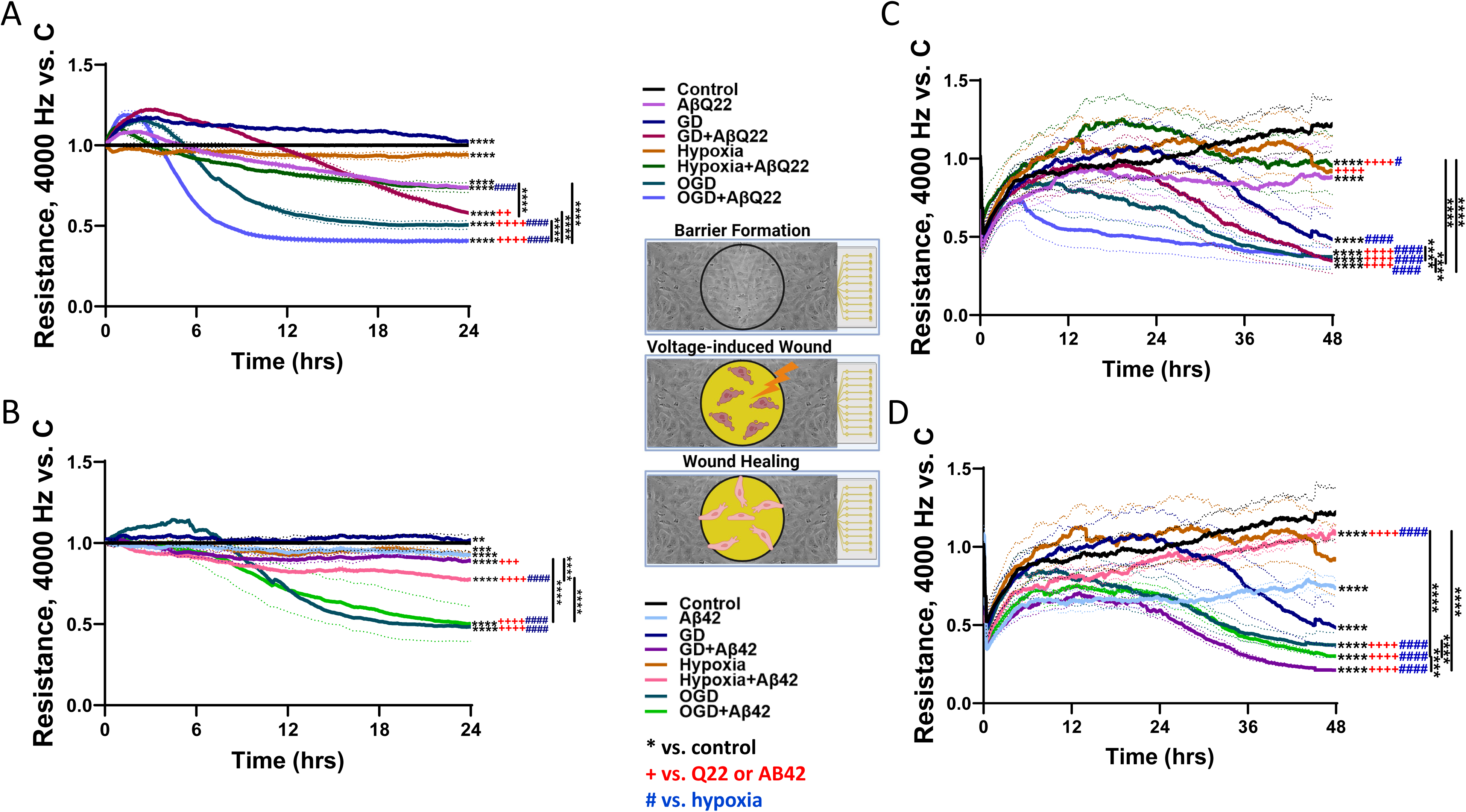
Potentiated impairments in TEER and wound healing during exposure of HCMECs to AβQ22 and Aβ42 challenge, under conditions of hypoxia, GD or OGD. **(A/B)** HCMECs were allowed to form a stable barrier for 48h and, post-barrier formation, were challenged with 25µM AβQ22 **(A)** or 5µM Aβ42 **(B)**, GD, or a combination of the two under conditions of normoxia and hypoxia. TEER was recorded for 24h post-treatment using the ECIS Zθ system (Applied Biophysics). Data are represented as Resistance change vs. normoxia control (black line). N=3 experiments with 2 technical replicates; a representative image from one experiment is shown; one-way ANOVA, Tukey’s post-test. **(C/D)** HCMECs were allowed to form a stable barrier for 48h and, post barrier formation, were treated with 25µM AβQ22 **(C)** or 5µM Aβ42 **(D)**, GD, or a combination of the two under conditions of normoxia and hypoxia. 1h post-treatment, cells were wounded at the center of the well for 20 seconds (60000hz). HCMEC TEER was then monitored for 48h post-wound and wound healing was indicated by post-wound TEER increases. Data is represented as resistance change vs. normoxia control (black line). N= 3 experiments with 2 technical replicates; one-way ANOVA, Tukey’s post-test (**** p<0.0001, ***p<0.001, **p<0.01, *p<0.05 vs. normoxia control) (**++++** p<0.0001, +++ p<0.001, ++ p<0.01, + p<0.05 vs. AβQ22 or Aβ42) (#### p<0.0001, ### p<0.001, ## p<0.01, # p<0.05 vs. hypoxia).

To assess wound repair ability, a wound healing assay was conducted utilizing a dedicated ECIS assay. Following barrier formation, HCMECs were treated with 25µM AβQ22 or 5µM Aβ42, GD, or a combination of both under normoxia or hypoxia conditions. 1h post-treatment, a strong electrical current was applied to the center of the EC barrier to inflict a circular wound. Recovery from the wound was monitored for 48h and was indicated by post-wound TEER increases. As our lab has recently demonstrated, AβQ22-treated HCMECs demonstrated impaired wound healing versus controls (**Fig. 2C**) [42]. Similarly, treatment of HCMECs with Aβ42 resulted in impaired wound healing versus controls (**Fig. 2D**). Combined exposure of HCMECs to AβQ22+GD exacerbated wound healing impairments compared to cells treated with each challenge alone, while HCMECs treated with AβQ22+OGD demonstrated an additive decrease in wound healing levels (**Fig. 2C**). Interestingly, Aβ42+OGD did not have an additive effect on HCMEC wound healing versus OGD alone, but HCMECs treated with Aβ42+GD, Aβ42+OGD, or OGD alone, all demonstrated the most impaired wound healing capabilities (**Fig. 2D**).

### Exposure of HCMECs to AβQ22 or Aβ42 in combination with OGD exacerbates changes in expression of BBB-modulating proteins

Strong evidence from previous *in vitro*, *in vivo*, and post-mortem human studies indicate that Aβ species can promote BBB permeability by disrupting the expression and phosphorylation state of BBB-regulating proteins, such as tight junction (TJ), immune cell extravasation proteins, and cadherins [43–47]. Additionally, exposure of brain ECs to OGD has previously been reported to disrupt the expression and regulation of barrier proteins [30, 48, 49]. Since Aβ and OGD both have been found to cause dysfunctional modulation of cerebral endothelial barrier-proteins, we aimed to understand whether combined exposure of HCMECs to Aβ+OGD additively exacerbates expressional changes within BBB-related proteins and whether these changes are Aβ-species specific and correlate to the TEER decreases observed in **Figure 2**. To do so, we analyzed zona occludin 1 (ZO1), phosphorylated claudin-5 (pClaudin5), and MMP2 (matrix metalloproteinase 2) protein expression in HCMECs treated with 25µM AβQ22 or 5µM Aβ42, GD, or a combination of both under normoxia or hypoxia conditions for 24h, a timepoint revealing drastic TEER decreases (**Figure 2)**. ZO1 is a vital anchor protein necessary for organizing TJ proteins and connecting them to the actin cytoskeleton [50]. Hypoxia-treated HCMECs demonstrated a significant increase in ZO1 expression versus controls, possibly indicating a HIF-1 mediated compensatory mechanism to maintain barrier integrity under low oxygen [51] **(Fig. 3A/B)**. However, HCMECs treated with AβQ22 under hypoxia or OGD conditions demonstrated significant decreases in ZO1 expression compared to hypoxia controls, with an additive decrease in ZO1 expression being revealed by HCMECs treated with AβQ22+OGD **(Fig. 3A)**. Similarly, HCMECs treated with Aβ42+OGD demonstrated a decrease in ZO1 expression compared to Aβ42 alone or Aβ42+hypoxia, but the decrease did not reach the same significance as in AβQ22+OGD-exposed HCMECs **(Fig. 3B)**. Claudin-5 is a major TJ protein, and its phosphorylation has been directly linked to increased endothelial barrier permeability [52]. Interestingly, we revealed that OGD challenge increased pClaudin5 expression versus controls. Moreover, HCMECs treated with AβQ22+OGD or Aβ42+OGD demonstrated an additive increase in pClaudin5 expression compared to cells challenged with the peptides alone, with Aβ42+OGD more significantly increasing pClaudin5 levels than AβQ22+OGD **(Fig. 3A/B)**.

**Figure 3.**
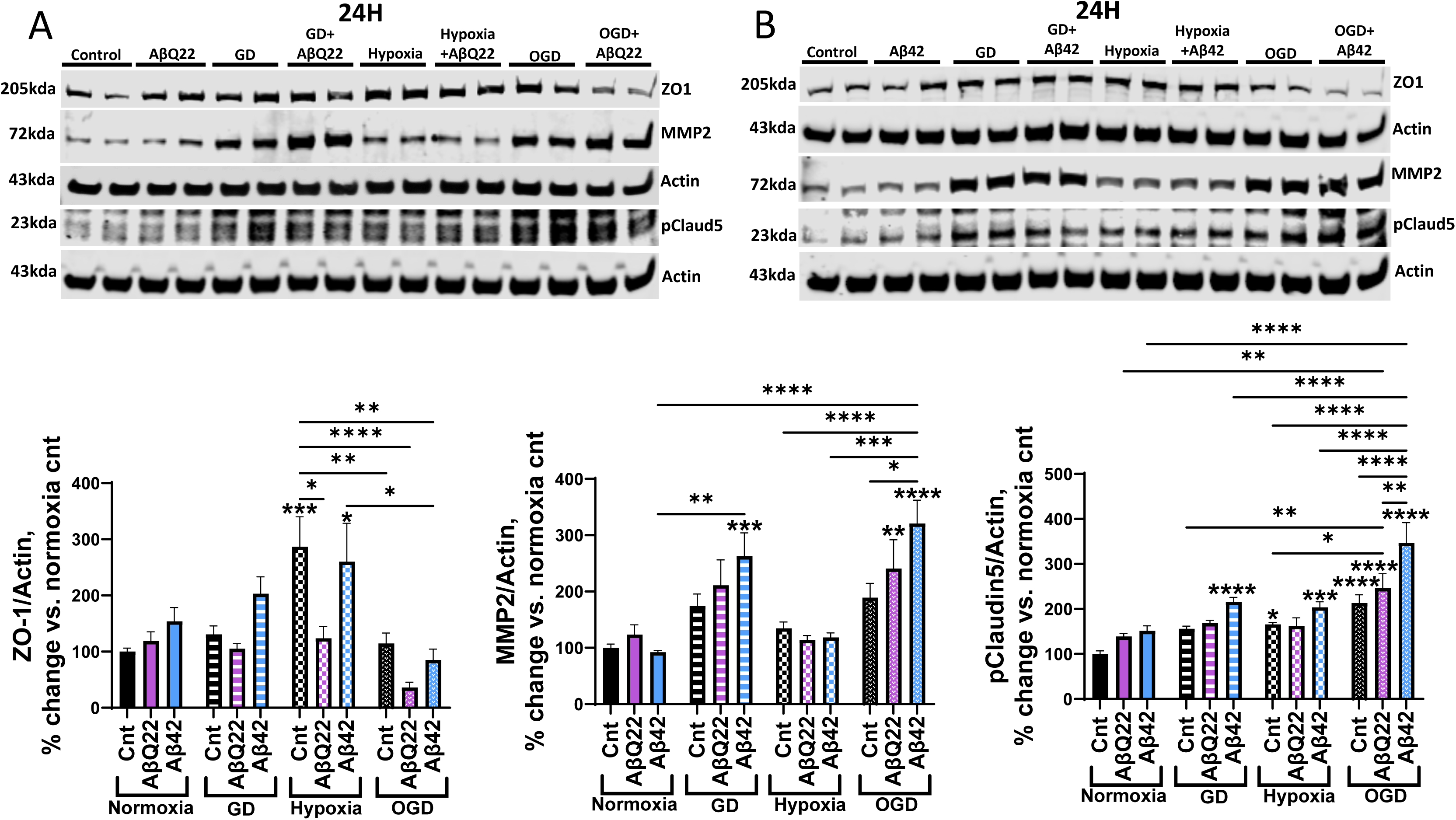
Expression changes in BBB-modulating proteins induced by treatment with AβQ22 or Aβ42 in the presence of GD, hypoxia or OGD. **(A/B)** HCMECs were treated with 25µM AβQ22 **(A)** or 5µM Aβ42 **(B)**, GD, or a combination of both under conditions of normoxia or hypoxia for 24h. ZO1, MMP2, and pClaudin5 protein expression was evaluated via WB and actin was used for normalization. Data is represented as % change vs. normoxia control. N=3 experiments with 2 technical replicates. two-way ANOVA, Tukey’s post-test; * located over bars are comparisons vs. normoxia control (**** p<0.0001, ***p<0.001, **p<0.01, *p<0.05).

MMP2 is an enzyme responsible for degrading the extracellular matrix and is a known player in BBB breakdown and cerebral hemorrhage in AD and CAA [53, 54]. Our lab and others have demonstrated that exposure of HCMECs to Aβ species, particularly AβQ22, increases MMP2 expression and activity [27, 42]. Similarly, OGD has been reported to induce upregulation of MMP2 expression in HCMECs and increase their MMP2 secretion [55]. Following 24h treatment, HCMECs challenged with AβQ22 or Aβ42 under conditions of GD and OGD demonstrated potentiated increases in MMP2 protein expression, with HCMECs treated with Aβ42+OGD revealing the most significant increase in MMP2 levels **(Fig. 3A/B)**. We also measured the expression of these proteins following an acute treatment (6h), a timepoint where TEER decreases began to be observed (**Figure 2)**. Very similar trends to the 24h timepoint were observed at this early timepoint **(Supplemental Fig. 1)**. Overall, the exacerbated ZO1 loss, claudin-5 phosphorylation, and MMP2 upregulation induced by exposure of HCMECs to Aβ+OGD indicate that Aβ and OGD act in an additive manner on the same protein mediators to potentiate the loss of blood-brain barrier (BBB) function.

### AβQ22 or Aβ42 combined with OGD differentially potentiate increases in cerebrovascular inflammatory modulators and monocyte trans-endothelial extravasation

Intercellular adhesion molecule 1 (ICAM1) is a key player in the extravasation of immune cells from the lumen of cerebral vessels into the brain parenchyma, where these immune cells release proinflammatory factors and drive neuroinflammation in AD and diseases associated with CH [56]. Additionally, proinflammatory cytokines, including IL-6, IL-8, and IFN-γ, have been shown to induce BBB permeability, promoting reduced TJ expression and improper TJ localization [57–59]. To understand whether combined exposure of HCMECs to Aβ species and OGD exacerbates the expression of proinflammatory mediators, we measured ICAM1 protein expression in HCMECs treated with 25 µM AβQ22 or 5 µM Aβ42, GD, or a combination of both for 24h under normoxia or hypoxia conditions. While GD alone resulted on an increase in ICAM expression, AβQ22+GD as well as AβQ22+OGD induced potentiated increases in ICAM1 expression. This overexpression appeared to be mainly driven by GD and not hypoxia **(Fig. 4A)**. Differentially from AβQ22, Aβ42 challenge did not cause additive effects on GD-mediated ICAM1 overexpression **(Fig. 4A)**.

**Figure 4.**
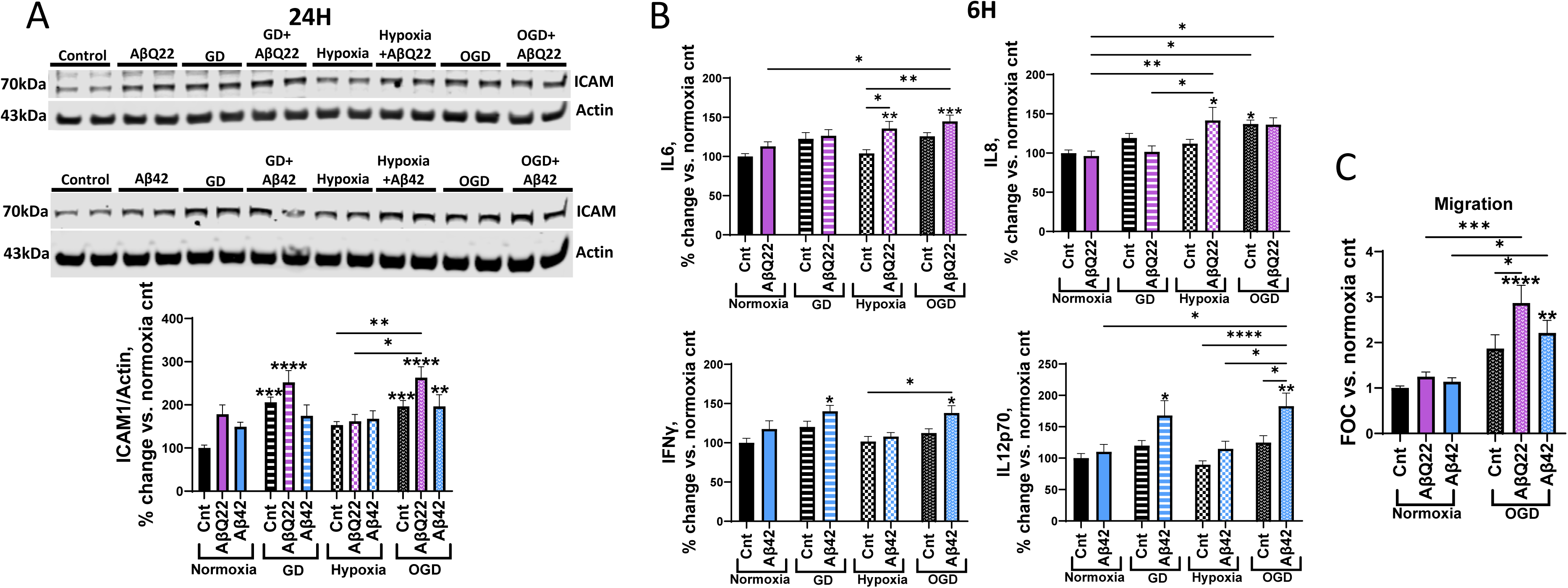
AβQ22 or Aβ42 under hypoxia, GD or OGD differentially effect ICAM1 expression, pro-inflammatory cytokine secretion in HCMECs, or Aβ-induced monocyte migration. **(A)** HCMECs were treated with 25µM AβQ22 **(A)** or 5µM Aβ42 **(C)**, GD, or a combination of both under conditions of normoxia or hypoxia for 24h. ICAM1 protein levels were evaluated via WB and actin was utilized for normalization. Data is represented as % change vs. normoxia control (N=3 experiments with 2 technical replicates; two-way ANOVA, Tukey’s post-test). **(B)** HCMECs were treated with 25µM AβQ22 or 5µM Aβ42, GD, or a combination of the two under conditions of normoxia or hypoxia for 6h. Media was collected to run a multiplex pro-inflammatory cytokine assay (MSD), and protein concentration was utilized for normalization. Data is represented as % change vs. normoxia control (N=3 experiments with 2 or more technical replicates; two-way ANOVA; Tukey’s post-test). **(C)** HCMECs were allowed to form confluent monolayers on trans-well inserts and were subsequently treated with 25µM AβQ22 or 5µM Aβ42, OGD, or a combination of the two for 24h. Following treatment, Calcein-AM labeled U937 monocytes were allowed to migrate across the EC barrier for 6h and fluorescence was measured to determine the number of migrated monocytes. Data is represented as FOC vs. normoxia control (N=3 experiments with 2 technical replicates; two-way ANOVA; Tukey’s post-test). * over bars are comparisons to the normoxia control (**** p<0.0001, ***p<0.001, **p<0.01, *p<0.05).

Furthermore, we aimed to understand whether combined exposure of HCMECs to Aβ species and OGD additively promoted the secretion of proinflammatory cytokines. To do so, we utilized a multiplex cytokine array (MSD). HCMECs were treated with 25µM AβQ22 or 5µM Aβ42, GD, or a combination of both under normoxia or hypoxia conditions for 6h, and media was collected. The 6h timepoint was selected to understand whether early changes in HCMEC inflammatory state contribute to the decreases in TEER observed in **Figure 2**. AβQ22 and Aβ42 differentially affected the expression of certain proinflammatory cytokines. Treatment of HCMECs with AβQ22+OGD resulted in an additive increase in IL6 secretion, a potent inducer of BBB permeability, versus controls and cells exposed to each treatment alone (**Fig. 4B**). HCMECs treated with AβQ22+OGD also demonstrated a significant increase in IL8 expression, another cytokine linked to decreased BBB integrity, compared to control cells, but this effect appeared to be similar also in cells challenged with hypoxia+AβQ22 and with OGD alone (**Fig. 4B**). Conversely, treatment of HCMECs with Aβ42 and GD or OGD resulted in a comparable potentiation in IFN-γ secretion levels, another cytokine that is known to promote BBB breakdown (**Fig. 4B**). Additionally, treatment of HCMECs with Aβ42+GD or Aβ42+OGD resulted in a significant increase in IL12p70, a pleotropic proinflammatory cytokine **(Fig. 4B)**. Other proinflammatory cytokines were measured and did not reveal significant differences between treatments (**Supplemental Fig. 2**).

To understand whether the combined challenge with Aβ+OGD also impacted a functional measure of the microvascular inflammatory activation, a monocyte trans-endothelial extravasation assay was conducted. HCMECs were allowed to form monolayers and were subsequently treated with 25µM AβQ22 or 5µM Aβ42, OGD, or a combination of both for 24h. Post-treatment, fluorescently labeled human monocytes were allowed to migrate across EC barriers for 6h and the number of extravasated monocytes was determined. Treatment of HCMECs with a combination of either AβQ22 or Aβ42 and OGD resulted in a significant potentiation of monocyte migration across EC monolayers, with HCMECs exposed to AβQ22+OGD demonstrating the most significant increase in monocyte migration versus controls and each treatment condition alone **(Fig. 4C)**.

Taken together, the increases in ICAM1 and proinflammatory cytokines expression resulting from HCMEC exposure to Aβ+OGD unveil specific molecular mechanisms through which Aβ+OGD work in concert to exacerbate EC activation, which results in increased monocyte migration across the HCMEC barrier, thus providing insights into the pathways promoting HCMEC barrier permeability and microvascular inflammatory activation in conditions with comorbid vascular amyloidosis and hypoperfusion.

### Aβ and OGD additively decrease HCMECs angiogenic capabilities

Cerebral angiogenesis is an essential homeostatic process allowing ECs to form new vessels to maintain cerebral perfusion levels needed for proper brain function, particularly in conditions of poor perfusion or microvascular damage [60]. Our lab has demonstrated that challenging HCMECs with low doses of AβQ22 and Aβ42 impairs angiogenesis, specifically decreasing vessel branching [26]. Similarly, previous evidence reveals that OGD-exposed ECs demonstrate impaired angiogenic capabilities, specifically decreased vessel sprouting, migration, and tube formation [34, 35]. Hence, we sought to elucidate whether exposure of HCMECs to Aβ in combination with OGD potentiates angiogenic impairment and whether both challenges operate through mutual molecular mechanisms to produce this dysfunction. HCMECs were treated with 1µM AβQ22 or Aβ42 (the lowest dose capable of inducing angiogenesis inhibition in previous studies [26, 28]), GD, or a combination of the two, and the ability to form vessels was monitored after 6h under normoxia or hypoxia, through an angiogenesis assay. HCMECs demonstrated significantly decreased vessel branch numbers versus controls in response to all treatment conditions **(Fig. 5A)**, indicating decreased angiogenesis. Specifically, either peptide alone, as well as GD or hypoxia alone, significantly inhibited angiogenesis. However, HCMECs exposed to Aβ peptides+OGD demonstrated the most drastic inhibition of vessel branching versus cells exposed to the treatments alone **(Fig. 5A)**.

**Figure 5.**
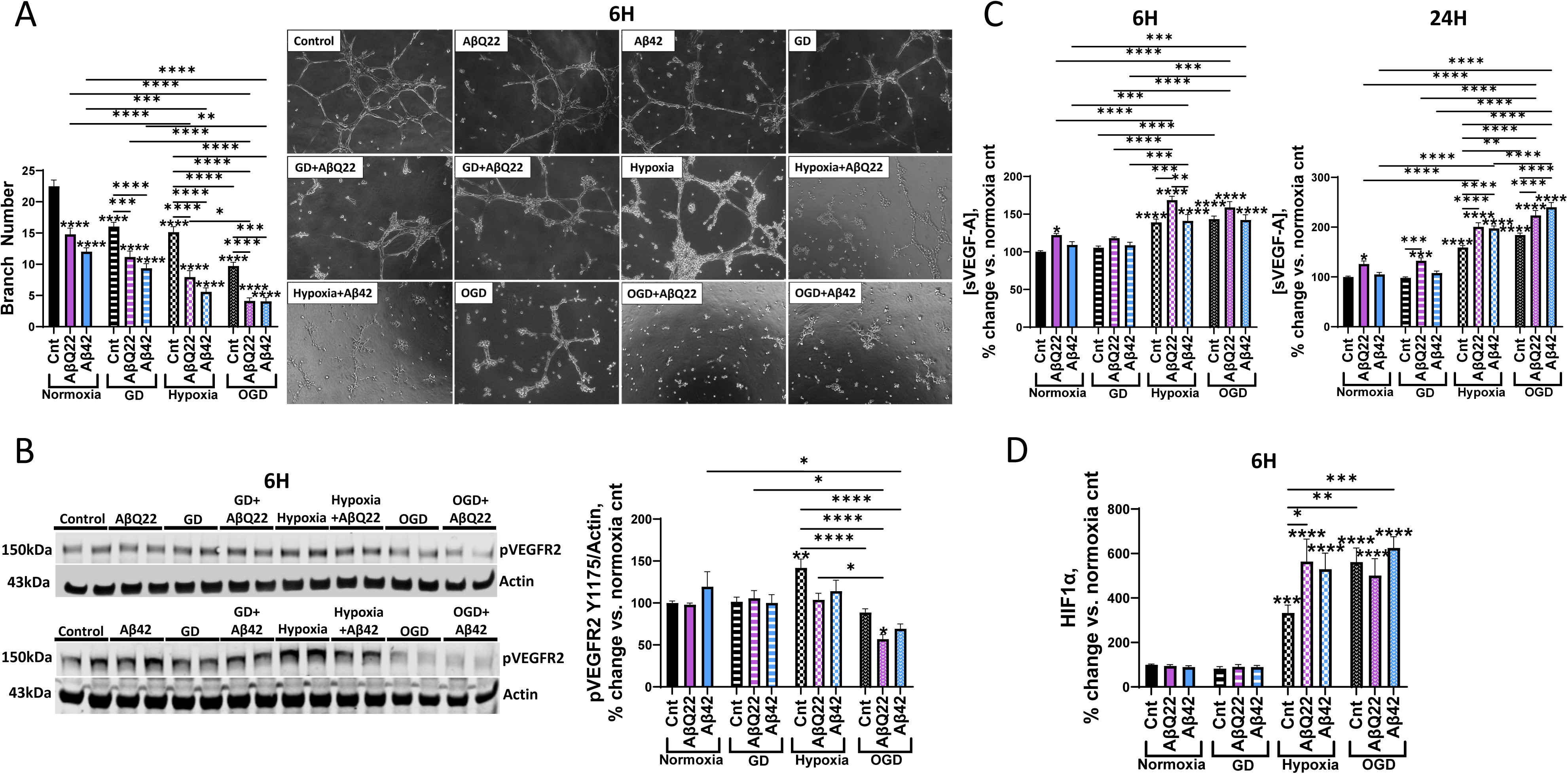
HCMECs exposed to AβQ22 or Aβ42 and OGD demonstrate potentiated impairment in angiogenic capabilities, despite increased sVEGF-A and HIF1. **(A)** HCMECs were treated with AβQ22 or Aβ42, GD, or a combination of both under conditions of normoxia or hypoxia for 6h. Vessel branch number was evaluated with an angiogenesis inhibition assay kit (Millipore). Data is represented as % change vs. normoxia control (N=3 experiments with three or more technical replicates; two-way ANOVA, Tukey’s post-test). **(B)** HCMECs were treated with AβQ22 or Aβ42, GD, or a combination of both under conditions of normoxia or hypoxia for 6h. pVEGFR2 protein levels were evaluated by WB and actin was used for normalization. Data is represented as % change vs. normoxia control (N=3 experiments with 2 technical replicates; two-way ANOVA, Tukey’s post-test). **(C)** HCMECs were treated with AβQ22 or Aβ42, GD, or a combination of the two under conditions of normoxia or hypoxia for 6h **(left)** and 24h **(right)**. Media was collected to conduct a sVEGF-A ELISA. Data is represented as % change vs. normoxia control (N=3 experiments with two technical replicates; two-way ANOVA, Tukey’s post-test). **(D)** HCMECs were treated with AβQ22 or Aβ42, GD, or a combination of the two under conditions of normoxia or hypoxia for 6h. Cell lysate was collected to run a HIF1α ELISA. Data is represented as % change vs. normoxia control (N=3 experiments with two technical replicates; two-way ANOVA, Tukey’s post-test). *s located over bars are comparisons vs. normoxia control (**** p<0.0001, ***p<0.001, **p<0.01, *p<0.05).

Since combined exposure of HCMECs to Aβ+OGD additively decreased angiogenic capabilities, we then aimed to determine whether we observed analog changes in the protein expression of key angiogenic mediators. HCMECs were treated with Aβ peptides, GD, or a combination of both at the same 6h timepoint under normoxia or hypoxia conditions. Protein expression of the phosphorylated vascular endothelial growth factor receptor 2 (VEGFR2) Y1175 (pVEGFR2), the VEGFR2 phosphorylated at tyrosine1175, a known phosphorylation site associated with angiogenic signaling activation, was evaluated. As expected, hypoxia-exposed HCMECs demonstrated a significant increase in pVEGFR2 expression versus controls **(Fig. 5B).** Hypoxic conditions are known to stabilize the transcription factor HIF1α, which transcribes VEGF. VEGF subsequently binds to the VEGFR2 to induce its phosphorylation and activation [61]. The hypoxia-induced increase in active pVEGFR2 was prevented when HCMECs were exposed to hypoxia+AβQ22 or Aβ42 **(Fig. 5B)**. HCMECs exposed to Aβ42+OGD demonstrated a decrease in pVEGFR2 compared to cells treated with Aβ42 alone, while cells challenged with AβQ22+OGD demonstrated the most significant decrease versus hypoxia and normoxia controls **(Fig. 5B)**. Accordingly, we found that HCMECs treated with AβQ22+OGD demonstrated a significant decrease in VEGF-A intracellular protein abundance versus hypoxia and normoxia controls **(Supplemental Fig. 3)**. Because VEGF-A is normally secreted by vascular ECs, we also measured soluble VEGF-A (sVEGF-A) levels within HCMEC-conditioned media after 6h and 24h. At both the acute (6h) and chronic (24h) timepoints a significant increase in sVEGF-A levels was observed when HCMECs were exposed to hypoxia, coinciding with previous studies **(Fig. 5C).** Interestingly, AβQ22-exposed HCMECs demonstrated a significant increase in sVEGF-A release versus controls at 6h and 24h **(Fig. 5C)**. Specifically, at 6h, HCMECs treated with AβQ22 under hypoxia conditions revealed the highest sVEGF-A level versus normoxia and hypoxia controls, while Aβ42 treatment had no effect on sVEGF-A expression **(Fig. 5C)**. At the 24h timepoint an additive increase in sVEGF-A level was observed when cells were exposed to Aβ42+OGD and the same additive increase was observed when HCMECs were treated with AβQ22+OGD **(Fig. 5C)**. Overall, these results suggest that an increased release of sVEGF-A from the cells may be a compensatory mechanism in response to the hypoxia/OGD. However, this does not result in a functional VEGFR2 activation, particularly in the combination treatment with Aβ and OGD.

HIF1α is the main regulator of the hypoxic response and the transcription factor that promotes pro-angiogenic gene expression, such as VEGF. Therefore, we also measured HIF1α protein expression in HCMECs treated with AβQ22 or Aβ42, GD, or a combination, under normoxia or hypoxia conditions for 6h. The 6h timepoint was chosen due to previous evidence demonstrating that 6h of hypoxia results in peak HIF1α expression in vascular ECs [62]. As anticipated, exposure of HCMECs to hypoxia alone resulted in a significant increase in HIF1α expression **(Fig. 5D)**. Treatment of HCMECs with AβQ22 under hypoxic conditions significantly potentiated HIF1α increases, but no further increase was observed from AβQ22+OGD treatment **(Fig. 5D)**. HCMECs treated with Aβ42+OGD demonstrated the most significant upregulation in HIF1α protein expression versus normoxia and hypoxia controls, which appeared mainly driven by the OGD treatment. Taken together, these data suggest that Aβ potentiates the hypoxic/OGD response, exacerbating HIF1α upregulation as well as sVEGF-A secretion, but these pro-angiogenic factors accumulate without promoting angiogenic signaling, since we observe an additive decrease in pVEGFR2 expression and vessel branching after Aβ+OGD challenge.

## Discussion

AD is a multifactorial syndrome not only driven by Aβ and tau accumulation. It is now well accepted that the vascular contributions to cognitive impairment and dementia (VCID), and particularly vascular disruptions resulting in CH are key drivers of early AD pathogenesis, even before considerable Aβ deposition [10–13, 63–65]. Indeed, since vascular and amyloid pathologies constitute the most frequent combination of mixed etiology dementias, Aβ accumulation can often occur within an already hypoperfused cerebral environment, where EC death, barrier dysfunction, and angiogenic impairment may be common consequences of the combined exposure to both Aβ and OGD. This study sought to determine whether exposure of HCMECs to Aβ and OGD results in additive increases in common molecular mediators of apoptotic and necrotic cell death mechanisms, barrier permeability, and angiogenesis failure. Additionally, we aimed to understand whether two different Aβ species that are known to be pro-apoptotic for cerebral ECs and to promote BBB permeability (AβQ22 and Aβ42 [26, 28, 66]) promote similar or differential mechanisms of HCMEC dysfunction in hypoxia, GD, and OGD conditions.

We began by investigating how the combined exposure of HCMECs to Aβ+OGD influences the activation of cell death mechanisms. Our lab has previously demonstrated that both AβQ22 and Aβ42 increase HCMEC apoptosis and, later, necrosis levels [4, 26, 67–69]. Hypoxia has also been linked to increased apoptosis and necrosis [70] and OGD *in vitro* has been found to specifically increase microvascular EC death through the same ischemia-mediated cell death pathways [29]. This study demonstrates that exposure of HCMECs to AβQ22 or Aβ42 in combination with GD significantly potentiates caspase-3 activation. Similarly, we observed a potentiated increase in apoptosis levels in HCMECs exposed to AβQ22+GD, while an additive increase in necrosis levels was observed in HCEMCs exposed to Aβ42+OGD, particularly after HR injury. Overall, these results suggest that lack of glucose, in the presence of AβQ22, pushes HCMECs towards potentiated apoptosis, while reduction and variability in oxygen levels, such as during HR, drives AβQ22-exposed HCMECs towards necrosis. Both AβQ22- and Aβ42-exposed HCMECs under GD conditions demonstrate potentiated increases in cleaved caspase-3, with this trend being significantly more evident with Aβ42 treatment. This strong caspase-3 activation at 24h in Aβ42-exposed cells may accelerate the terminal stages of apoptosis, shifting HCMEC fate to secondary necrosis faster, which was particularly evident in HCMECs treated with Aβ42+OGD. These results also confirm that manipulation of oxygen levels, particularly during HR, is sufficient to potentiate necrotic cell death in cerebral ECs, even in the absence of Aβ peptides. In line with these findings, it is known that in ischemic stroke, where oxygen delivery is disrupted, necrosis is the major cell death mechanism within the ischemic core and ischemia/reperfusion injury is highly associated with necrosis, with this tissue necrosis often resulting in substantial activation of neuroinflammatory processes [71–73].

Cerebral ECs ability to preserve proper barrier integrity is crucial for maintenance of healthy BBB structure and function. Increased BBB permeability is a hallmark of AD pathology, with a collection of studies reporting leakage of blood proteins as well as loss of endothelial, basement membrane, and TJ proteins within post-mortem AD brains [74]. Prior studies from our lab have demonstrated that Aβ species (AβQ22/Aβ42) are capable of decreasing TEER, an indicator of EC barrier permeability [26] [28]. Similar to AD, ischemic strokes are also known to cause substantial BBB breakdown, resulting from dysregulation of TJs and upregulation of proinflammatory mediators and MMP activity [75]. Through this study, we provide novel evidence that combined exposure of HCMECs, specifically to AβQ22+OGD, results in a continued additive decrease in TEER, reflecting the most permeable HCMEC barrier. This additive effect also appears at early (before 12h), but not late time points for Aβ42+OGD, suggesting that the combined treatment may reach maximal toxicity early after challenge.

To dissect the molecular players involved in promoting the drastic EC barrier permeability displayed by Aβ+OGD-challenged HCMECs, we measured the expression of proteins that mediate EC barrier integrity. We found that ZO1, a BBB anchor protein that links TJs and cytoskeletal proteins, was significantly upregulated in response to hypoxia. Similarly, a recent study utilizing a multi-cellular cerebral organoid found a significant increase in vascular ZO1 expression following hypoxia exposure [51]. This hypoxia-induced ZO1 upregulation could reflect a possible compensatory mechanism employed by HCMECs to contrast barrier damage upon oxygen deprivation, but the structure and localization of this upregulated ZO1 would need to be explored to understand if this modulation is functional. Despite this, when HCMECs are challenged with AβQ22 or Aβ42 together with OGD, ZO1 expression is significantly decreased versus hypoxia controls, with specifically AβQ22+OGD-challenged cells presenting decreased ZO1 expression versus untreated controls **(Supplemental Fig. 4)**. If ZO1 upregulation is a compensatory mechanism for cerebral EC barrier maintenance under hypoxia and this upregulation is blocked when the additional Aβ challenge is introduced, this could explain why HCMECs exposed to hypoxia alone maintain TEER in close proximity to control cells, while HCMECs exposed to hypoxia/OGD + Aβ demonstrate significantly reduced TEER. Additionally, we found that HCMECs exposed to Aβ+OGD demonstrated the most significant increase in pClaudin5 expression. Claudin-5 phosphorylation is a post-translational modification associated with TJ instability and decreased EC barrier integrity, providing additional insight into possible molecular players involved in Aβ+OGD-induced TEER loss [52]. Interestingly, we also found that MMP2 expression was most significantly increased in HCMECs exposed to Aβ+OGD. MMP2, a matrix metalloprotease responsible for extracellular matrix breakdown, plays a major role in BBB dysfunction in AD and several Aβ species have been found to increase MMP2 expression and activity within cerebral ECs [27]. Moreover, activation of MMPs whose expression is HIF1α-dependent, including MMP2, is a characteristic molecular event within ischemic strokes [75]. Overall, the increase in MMP2 and pClaudin5 expression and decrease in ZO1 expression resulting from combined exposure of HCMECs to Aβ+OGD reveals BBB modulators that contribute to Aβ+OGD’s potentiated TEER loss.

It is known that the AD brain presents a chronic neuroinflammatory state, which promotes neurodegeneration and potentiates Aβ-associated pathology [76]. CH also triggers substantial neuroinflammatory events in efforts to restore the injured cerebral area, but in turn this also results in BBB breakdown, neurodegeneration, and cognitive impairment [77]. Here we have demonstrated that combined exposure of HCMECs to specifically AβQ22 and GD or OGD resulted in the potentiated expression of ICAM1, an immune cell adhesion molecule highly involved in promoting immune cell extravasation across the BBB. We also found differing inflammatory cytokines whose expression was affected upon AβQ22 or Aβ42 exposure in conditions of hypoxia or OGD. Both IL6 and IL8 secretion were significantly increased by HCMECs exposed to AβQ22+ hypoxia or +OGD. Interestingly, IL8 release was also increased by HCMECs exposed to OGD alone. Both IL6 and IL8 have been shown to directly cause cerebrovascular damage and BBB breakdown [57, 58]. Since AβQ22, the Dutch mutant, is strongly associated to CAA, cerebrovascular dysfunction, strokes and hemorrhages in the human familial disease,, its greater effect found here on potentiating vascular factors that promote inflammation and increasing secretion of cytokines associated with BBB permeability versus Aβ42, a parenchymal species, is highly correlated to the effects observed in the human pathology and can explain its exacerbated vascular toxicity.

Conversely, HCMECs treated with Aβ42 and GD or OGD demonstrated potentiated increases in IFN-γ and IL12p70. IL12p70, the biologically active heterodimeric form of IL12, mainly functions to promote IFN-γ production, and IFN-γ, in return, promotes more IL12 subunit production to continue the feedback loop, which is likely why both cytokines display similar trends within the same treatment condition [78]. IFN-γ has also been found to disrupt cerebral EC barriers, decreasing TEER and promoting trans-endothelial immune cell migration [59]. It is important to note that within the AD and CAA brain, Aβ40 and Aβ42 aggregated species coexist simultaneously on the vessel wall, thus they may concurrently promote the secretion of these differing cytokines, which may all work together to promote similar cerebrovascular damage and BBB dysfunction. Overall, our data underscores that exposure of HCMECs to Aβ+OGD potentiates a proinflammatory activation state within cerebral ECs and a leaky BBB. This was further confirmed by the observed increase in monocyte trans-endothelial migration, demonstrating that the inflammatory response is functional, with Aβ+OGD (particularly AβQ22+OGD) inducing exacerbated monocyte migration across the EC barrier.

Angiogenesis is another important homeostatic mechanism that is disrupted within AD. When CBF is reduced, like in AD or ischemic stroke, formation of new cerebral vessels is vital to restore proper brain perfusion. Increasing evidence demonstrates that aberrant VEGF/VEGFR2 signaling dysregulation may contribute to AD pathogenesis [79]. Our lab has demonstrated that Aβ species (AβQ22/Aβ42) potently inhibit HCMEC angiogenic capabilities [26, 28]. We observed that OGD blocked practically all wound healing ability of cerebral ECs, with AβQ22+OGD showing even more dramatic effects. HCMECs treated with both AβQ22 or Aβ42 and OGD also demonstrated the most impaired angiogenesis, as evidenced by the highest reduction in vessel branch number versus controls.

We showed that the reduction in phosphorylated (active) VEGFR2, pVEGFR2 Y1175, appears to be the molecular mediator that contributes to this angiogenic failure. When bound and auto-phosphorylated at tyrosine1175, pVEGFR2 activates downstream pro-angiogenic signaling to promote EC proliferation, migration, and new vessel formation [80]. We found a significant decrease in pVEGFR2 Y1175 expression in HCMECs treated with Aβ+OGD, with specifically AβQ22 revealing a significant decrease from controls as well as hypoxia-treated cells (which had, as expected, elevated VEGF and pVEGFR2 levels [61]). VEGF-A is the main ligand for VEGFR2, and HIF1α, a transcription factor stabilized under hypoxia, promotes transcription of pro-angiogenic genes, such as VEGF [81]. Interestingly, HCMECs exposed to AβQ22 or Aβ42 and OGD exhibited an exacerbated increase in sVEGF-A release, while the intracellular VEGF protein level was decreased, possibly due to the ECs more robustly releasing this growth factor into the media. This study is also the first to reveal that exposure of HCMECs to Aβ in a hypoxic environment potentiates HIF1α upregulation versus hypoxia alone. Overall, we can then hypothesize that exposure of cerebral ECs to Aβ within a hypoxic or ischemic environment exacerbates HIF1α overexpression, leading to increased transcription and overexpression of VEGF-A, which is released and should bind and activate VEGFR2. However, the observed decrease in pVEGFR2 in Aβ+OGD treated cells suggests that the VEGF-A overexpression is maladaptive, with VEGF-A not binding to and activating VEGFR2, thus leading to angiogenic failure. In line with these findings, VEGF has also been reported to be overexpressed within certain regions of the AD human brain [82] [18], and increased in human AD plasma [83], yet AD brains exhibit decreased microvessel density and a chronic state of CH and CBF decrease [79].

Although HCMECs exposed to AβQ22 and Aβ42 all demonstrated multiple mechanisms of EC impairment, this study also revealed subsets of EC dysfunction and specific molecular mediators that were differentially affected by specific Aβ species. HCMECs exposed to AβQ22 and varying glucose/oxygen conditions more strongly displayed increases in apoptosis, ICAM expression, and monocyte migration and decreases in TEER, ZO1 expression, and wound healing capabilities. In contrast, HCMECs exposed to Aβ42 and varying glucose/oxygen conditions more potently revealed exacerbated increases in necrosis, pClaudin5, and MMP2 levels. Additionally, AβQ22 and Aβ42 promoted increases in different cytokines, although all linked to EC barrier dysfunction. Overall, both AβQ22 and Aβ42 potentiated activation of cerebral EC death pathways as well as angiogenic impairments and EC barrier dysfunction, but AβQ22 appeared to exacerbate factors more strongly reflective of EC apoptosis, activation and barrier permeability, particularly in hypoperfusion conditions, directly correlating with functional increases in barrier permeability and inflammation.

Another interesting distinction that arose from this study was that different cerebral EC dysfunction mechanisms were produced in the face of GD versus hypoxia. GD had a greater influence on caspase-3 activation and apoptosis, while reduction in oxygen levels (hypoxia/HR) pushed HCMECs towards a necrotic fate. Interestingly, GD and hypoxia individually had only moderate effects on barrier integrity and wound healing abilities, but OGD caused dramatic defects in barrier and wound healing properties. We also showed that GD particularly exacerbates increases in MMP2 and ICAM expression. As previously discussed, we observed a significant increase in ZO1 expression in HCMECs exposed to hypoxia alone. Recent evidence suggests that brain microvascular ECs can form mature EC monolayers under hypoxic conditions and evidence of TJ protein upregulation, such as ZO1, under hypoxic conditions has been reported [84, 85]. Additionally, the HIF1α signaling pathway, which is upregulated during hypoxia, has been linked to elevated TJ protein expression [84]. As expected, under hypoxic conditions, HCMECs also displayed a notable HIF1α-mediated increase in pro-angiogenic factors (sVEGF-A, and pVEGFR2 Y1175), which is in line with the characteristic hypoxic response [61]. OGD potentiated the effects of hypoxia on decreasing TEER, wound healing, and vessel branching, and increasing pClaudin5, sVEGF-A, and HIF1α expression. However, OGD mitigated the hypoxia-induced increase in ZO1 and pVEGFR2, therefore resulting in even more impaired endothelial barrier function and angiogenesis.

One limitation of this study is that OGD *in vitro* does not directly mimic how CH operates in a living system, where cerebral ECs are exposed to other factors (changes in flow, shear stress, reperfusion following occlusion release) and other cells of the neurovascular unit, which would all also contribute to vessel damage and cerebral EC dysfunction in different ways. Despite this, this study’s experimental setting allowed us to mimic the specific endothelial effects of a nutrient/oxygen-deficient environment that would be expected within hypoperfused brain areas and allowed us to dissect whether deprivation of either oxygen, glucose, or their combination produced particular patterns of cerebral EC dysfunction, and elucidate their molecular mediators. Future studies should conduct similar experiments on microfluidic chips capable of maintaining flow in real time, or with disturbed flow. Additionally, future studies in progress in our lab and others are aiming to confirm these results in *in vivo* models such as a combined AD model with partial cerebral artery occlusions or cardiovascular risk factors, to clarify how diseases that induce chronic CH impact the severity of AD pathology and cognitive dysfunction.

### Conclusions

CBF impairment resulting in chronic CH is one of the earliest and most long-lasting pathological manifestations of AD and vascular dementia. Early in disease progression, hypoperfused cortical regions begin to display increased neuroinflammation, cell death, and BBB dysfunction, an environment perfectly primed to allow for poor clearance and increased Aβ accumulation [65, 86], thus exacerbating Aβ detrimental effects. Within this study, we have demonstrated that depriving HCMECs of oxygen and/or glucose potentiates Aβ-induced cerebral EC dysfunction, specifically promoting increased apoptosis and necrosis, barrier instability and dysregulation of BBB proteins, inflammatory activation, and angiogenesis and wound healing failure. This study also reveals important distinctions between the effects of two Aβ species that are known to impair brain EC function, and between GD and hypoxia. This work reveals specific molecular changes responsible for the detrimental effects of Aβ on HCMECs within an environment modeling CH and directly demonstrates that OGD renders HCEMCs more vulnerable to Aβ effects. As CBF disruptions and CH appear early during the progression of AD, this likely weakens the defenses of ECs lining cerebral vessels, resulting in potentiated detrimental effects of Aβ, worsened vascular dysfunction, loss of microvessels and additional CH, thus instigating a vicious cycle. Monitoring for cerebral perfusion abnormalities, especially in midlife individuals with higher dementia risk, could be a useful tool to control the risk of AD and dementia pathology or to possibly prevent or reduce disease severity and progression. Additionally, this work clarifies specific molecular mechanisms that are mutually activated by Aβ and OGD, revealing novel targets for the development of treatments, as well as possible early biomarkers for AD, CAA, vascular dementias, and particularly mixed vascular/AD dementias.

## Non-standard Abbreviations and Acronyms

AD: Alzheimer’s Disease
Aβ: Amyloid-beta
AβQ22: Aβ40E_22Q_
BBB: Blood brain barrier
CAA: Cerebral amyloid angiopathy
CBF: Cerebral blood flow
CH: Cerebral hypoperfusion
EC: Endothelial cell
FBS: Fetal bovine serum
GD: Glucose deprivation
HCMECs: Human cerebral microvascular endothelial cells
HR: Hypoxia reoxygenation
ICAM1: Intercellular adhesion molecule 1
LDH: Lactate dehydrogenase
MMP2: Matrix metalloproteinase 2
OGD: Oxygen glucose deprivation
pClaudin5: Phosphorylated claudin-5
pVEGFR2: Phosphorylated vascular endothelial growth factor receptor 2
sVEGF-A: Soluble vascular endothelial growth factor A
TEER: Trans-endothelial electrical resistance
TJ: Tight junction
VEGFR2: Vascular endothelial growth factor receptor 2
WB: Western blot
WT: Wildtype
ZO1: Zona occludin-1

## Acknowledgments

AC and SF designed the study. AC conducted the majority of experiments and wrote the manuscript draft. TB conducted the migration assays. SF oversaw the study, assisted with experimental planning, critically edited the manuscript, and obtained funding to support this project. All authors read and approved the final manuscript.

## Figure Legends

**Supplemental Figure 1.**
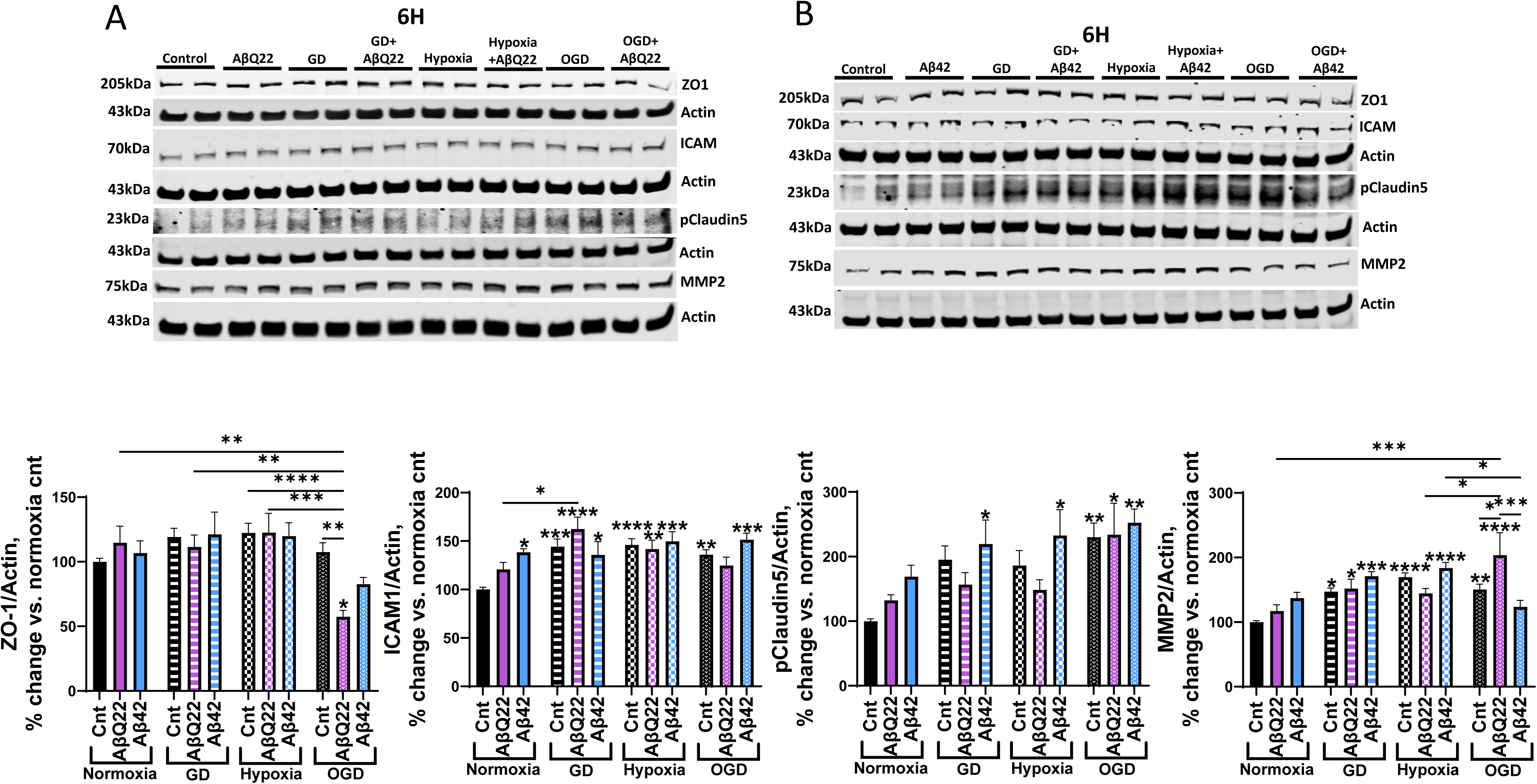
Modulation of BBB-related protein expression in HCMECs following acute treatment with AβQ22, Aβ42, and OGD. **(A/B)** HCMECs were treated with 25µM AβQ22 **(A)** or 5µM Aβ42 **(B)**, GD, or a combination of both under conditions of normoxia or hypoxia for 6h. ZO1, ICAM1, pClaudin5, and MMP2 protein expression was evaluated via WB and actin was used for normalization. Data is represented as % change vs. normoxia control. (N=3 experiments with 2 technical replicates. two-way ANOVA, Tukey’s post-test). * located over bars are comparisons vs. normoxia control (**** p<0.0001, ***p<0.001, **p<0.01, *p<0.05).

**Supplemental Figure 2.**
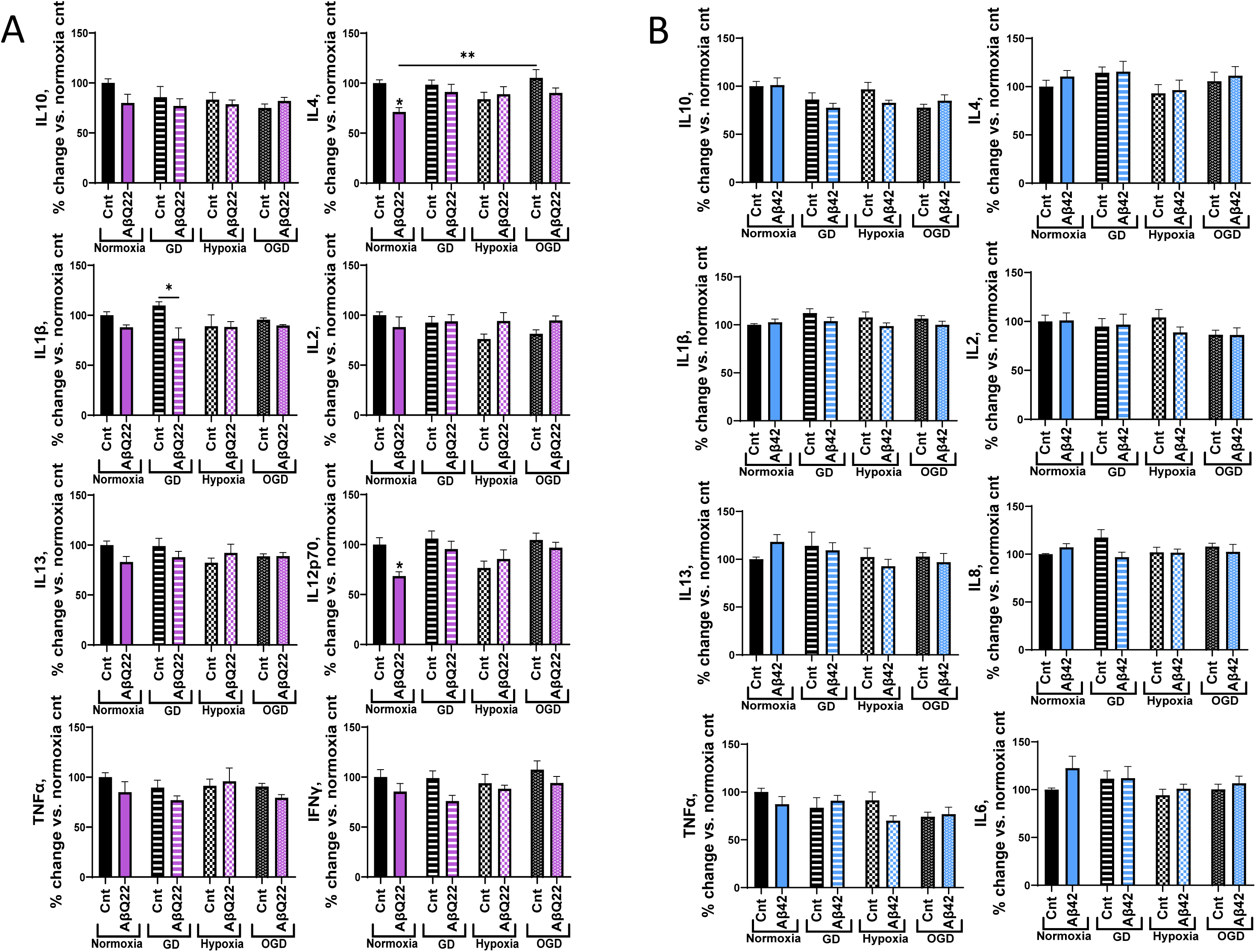
Effects of AβQ22, Aβ42, and OGD treatment on HCMEC release of additional pro-inflammatory cytokines. **(A/B)** HCMECs were treated with 25µM AβQ22 **(A)** or 5µM Aβ42 **(B)**, GD, or a combination of both under conditions of normoxia or hypoxia for 6h. Media was collected and utilized to run a multiplex pro-inflammatory cytokine assay (MSD). Protein concentration was utilized for sample normalization. Data is represented as % change vs. normoxia control (N=3 experiments with 2 technical replicates; one-way ANOVA, Tukey’s post-test). *s located over bars are comparisons vs. normoxia control. (**** p<0.0001, ***p<0.001, **p<0.01, *p<0.05).

**Supplemental Figure 3.**
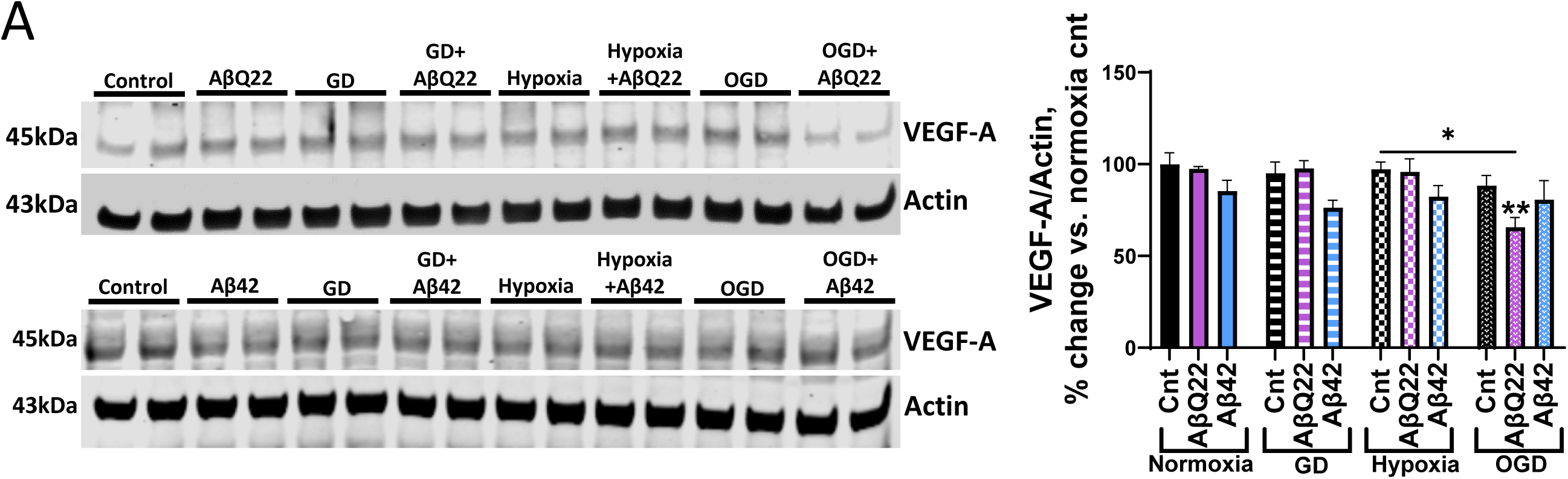
VEGF-A expressional changes following treatment with AβQ22, Aβ42, and OGD. **(A)** HCMECs were treated with 25µM AβQ22 or 5µM Aβ42, GD, or a combination of both under conditions of normoxia or hypoxia for 6h. VEGF-A protein expression was evaluated via WB analysis and actin was used for normalization. Data is represented as % change vs. normoxia control. N=3 experiments with 2 technical replicates. two-way ANOVA, Tukey’s post-test. * located over bars are comparisons vs. normoxia control (**** p<0.0001, ***p<0.001, **p<0.01, *p<0.05).

**Supplemental Figure 4.**
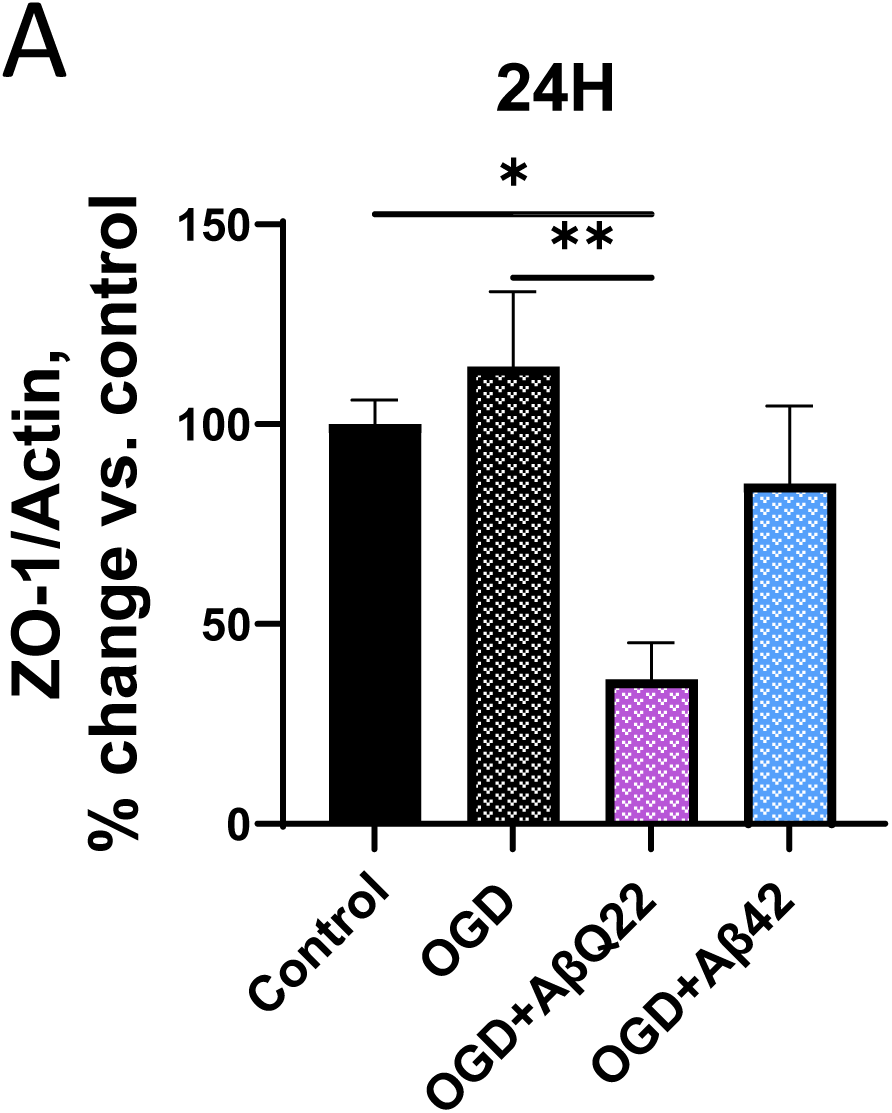
HCMECs treated with AβQ22+OGD reveal significantly decreased ZO1 protein expression compared to untreated controls as well as cells exposed to OGD alone. **(A)** 24h ZO1 protein expression data from Figure 3 comparing only OGD treatment groups to untreated control cells. Data is represented as % change vs. normoxia control (N=3 experiments with 2 technical replicates; one-way ANOVA, Tukey’s post-test). (**** p<0.0001, ***p<0.001, **p<0.01, *p<0.05.

